# Sex differences in vocalizations to familiar or unfamiliar females in mice

**DOI:** 10.1101/2020.06.13.150102

**Authors:** Eri Sasaki, Yuiri Tomita, Kouta Kanno

## Abstract

Mice, both wild and laboratory strains, emit ultrasound to communicate. The sex differences between male to female (male–female) and female to female (female– female) ultrasonic vocalizations (USVs) have been discussed for decades. In the present study, we compared the number of USVs emitted to familiar and unfamiliar females by both males (male–female USVs) and females (female–female USVs). We found that females vocalized more to unfamiliar than to familiar females. In contrast, males exhibited more USVs to familiar partners. This sexually dimorphic behavior suggests that mice change their vocal behavior in response to the social context, and their perception of the context is based on social cognition and memory. In addition, because males vocalized more to familiar females, USVs appear to be not just a response to novel things or individuals, but also to be a social response.

## 1. Introduction

Social animals utilize signals of specific modalities to communicate with other individuals. Among such signals, vocal communication is widely observed in animals and is superior in many respects (Brudzynski, 2010). Recently, vocal communication in mice has attracted considerable academic attention because of its power and utility in the investigation of molecular and neural mechanisms for social behaviors and their deficits, especially focusing on developmental disorders such as autism spectrum disorders (Fischer and Hammerschmidt, 2011; Lahvis et al., 2011; Konopka and Roberts, 2016). In addition, we previously investigated synapses and behaviors in the mutants of *Autism susceptibility candidate 2* (*AUTS2*) gene, and we reported that excitatory synaptic inputs were increased in the forebrain and that social interaction and vocalizations were altered in the mice (Hori et al., 2020). In mice, ultrasonic vocalizations (USVs) are used for communication, and they can be widely observed in both wild and laboratory strains (Sales, 2010). There are two types of USVs commonly known in mice: pupUSVs and courtship vocalizations (Konopka and Roberts, 2016). Mice pups emit ultrasounds (pupUSVs), mainly when they are outside their nest in the first several days after birth. A state of arousal due to the perception of cold; perception of unusual tactile stimulation; or loss of social contact triggers the emission of pupUSVs, which enhances the mother’s approach to the sound source (Ehret, 2005). The mother’s attention and movement leads to the expression of maternal behavior. On the other hand, adult male mice emit ultrasounds in the presence of adult females or their urine (Nyby et al., 1979); these are known as courtship vocalizations or courtship songs (Holy and Guo, 2005). Activational effects of sex hormones on male vocalizations have been reported (Nyby et al., 1992; Sipos and Nyby, 1996a). Furthermore, the number of male USVs is enhanced by sociosexual experience with females (Kanno and Kikusui, 2018), and usage of syllable types (call patterns) is modulated according to the progressive phase of sexual behaviors (Matsumoto and Okanoya, 2016). Thus, USVs from males to females are thought to be male-specific precopulatory behaviors (Burns-Cusato et al., 2004; Kanno and Kikusui, 2018). In addition, several studies have revealed that male USVs in turn affect female behaviors. Females of Swiss-Webster mice spent more time with intact vocalizing males than with devocalized males in the preference test. This preference could be mimicked by playback sounds and was not observed in ovariectomized females (Pomerantz et al., 1983). Playback of male vocalizations in B6 strain also elicited approach behaviors in females (Hammerschmidt et al., 2009), and playback of male USVs activated kisspeptin neurons, which play important roles in reproduction, as indicated by phosphorylated cyclic AMP response element binding protein (pCREB) immunoreactivity (Asaba et al., 2017). Moreover, females of B6 and BALB/c strains preferentially approach the playback vocalizations of the opposite strain (Asaba et al., 2014), suggesting that females can discriminate acoustic features of vocalization to a certain extent.

In this regard, vocalizations in mice involved in these two contexts have been well investigated since they were first reported approximately fifty years ago (Zippelius and Schleidt, 1956; Noirot, 1966; Sewell, 1967). Adult females were initially not thought to vocalize either to males or females (Nyby et al., 1977). However, using several strains, Maggio and Whitney meticulously revealed that adult females emit USVs to females (female–female), whereas they rarely emit to males (female–male) and that the number of female–female USVs is comparable to male–female courtship USVs (Maggio and Whitney, 1985). Thereafter, a study observing female–female USVs in albino NMRI female mice indicated that females vocalize more to unfamiliar females than to familiar ones (D’Amato and Moles, 2001). This finding suggests that female–female USVs reflect social cognition and memory. In fact, administration of a classical cholinergic antagonist (scopolamine) disrupted this social memory (D’Amato and Moles, 2001). Although female–female USVs are known to be related to this social memory, the functions of these USVs are poorly understood.

In the study by D’Amat and Moles (D’Amato and Moles, 2001) described above, familiar individuals were those that were previously presented only several minutes in advance (while simultaneously recording ultrasound). Therefore, we decided to use female individuals that were reared in the same cage after weaning as familiar individuals. In other words, we hypothesized that female–female USVs reflect social affiliation if females emit USVs more to familiar females than to unfamiliar ones under such conditions. If this hypothesis is correct, the results of the study by D’Amat and Moles and our expected results could be explained as follows: females express high levels of USVs to establish an affiliative relationship with a stranger of the same sex first; and, at the same time, females also emit more USVs to familiar social members. In the D’Amat and Moles study, 15-min, 30-min, 60-min, and 24-hour time intervals after female subjects encountered females that would be used as familiar individuals was applied. Subject females emitted more USVs to unfamiliar than to familiar females under three conditions of time intervals (15-min, 30-min, and 60-min), while the difference was not observed after the 24-hour interval. The effect of the cholinergic antagonist mentioned above was investigated at 30-min intervals because it represented the best interval for drug absorption. Thus, in this study, a 30-min interval that seemed suitable for retention time with versatile use and a 24-hour interval that seemed severe for mice retention, was applied to B6 mice.

In the present study, we also compared female–female vocalizations and male– female vocalizations using the B6 strain, which is currently the most commonly used laboratory strain. Social cognition and memory in male mice has not yet been measured in this context; however, it is important to measure the sociality of both sexes using the same methods because mouse models and USVs have been recently investigated to clarify the basic mechanisms of developmental disorders (Fischer and Hammerschmidt, 2011; Lahvis et al., 2011; Konopka and Roberts, 2016).

## 2. Materials and Methods

### 2.1 Animals

We used an inbred strain of B6 (C57BL/6J) mice. Mice were purchased from Japan SLC (Hamamatsu, Japan). Subject animals were used when they were 8–10 weeks old for the recording tests. All mice, after being purchased were reared in the same breeding room. The food (5L37 Rodent LabDiet EQ, PMI Nutrition International, MO, USA) and bedding (soft woodchip, Japan SLC, Hamamatsu, Japan) were sold by the same supplier, Japan SLC.

For recordings of male–female USVs, a male and an age-matched unfamiliar female mouse were cohoused in a standard cage (182 × 260 × 128 mm, CREA Japan) immediately following arrival from the supplier until experiments began. Pups were delivered in 4 of the 29 male–female pairs. Owing to such small N values, observation of delivery was not taken into account for analysis.

For recordings of female–female USVs, females that were bred together in the same cage after weaning were purchased, and two individuals were housed in each cage immediately after arrival from the supplier until experiments began. One was used as the subject, and the other was used as the familiar partner (see Fig. 1A).

**Fig. 1.**
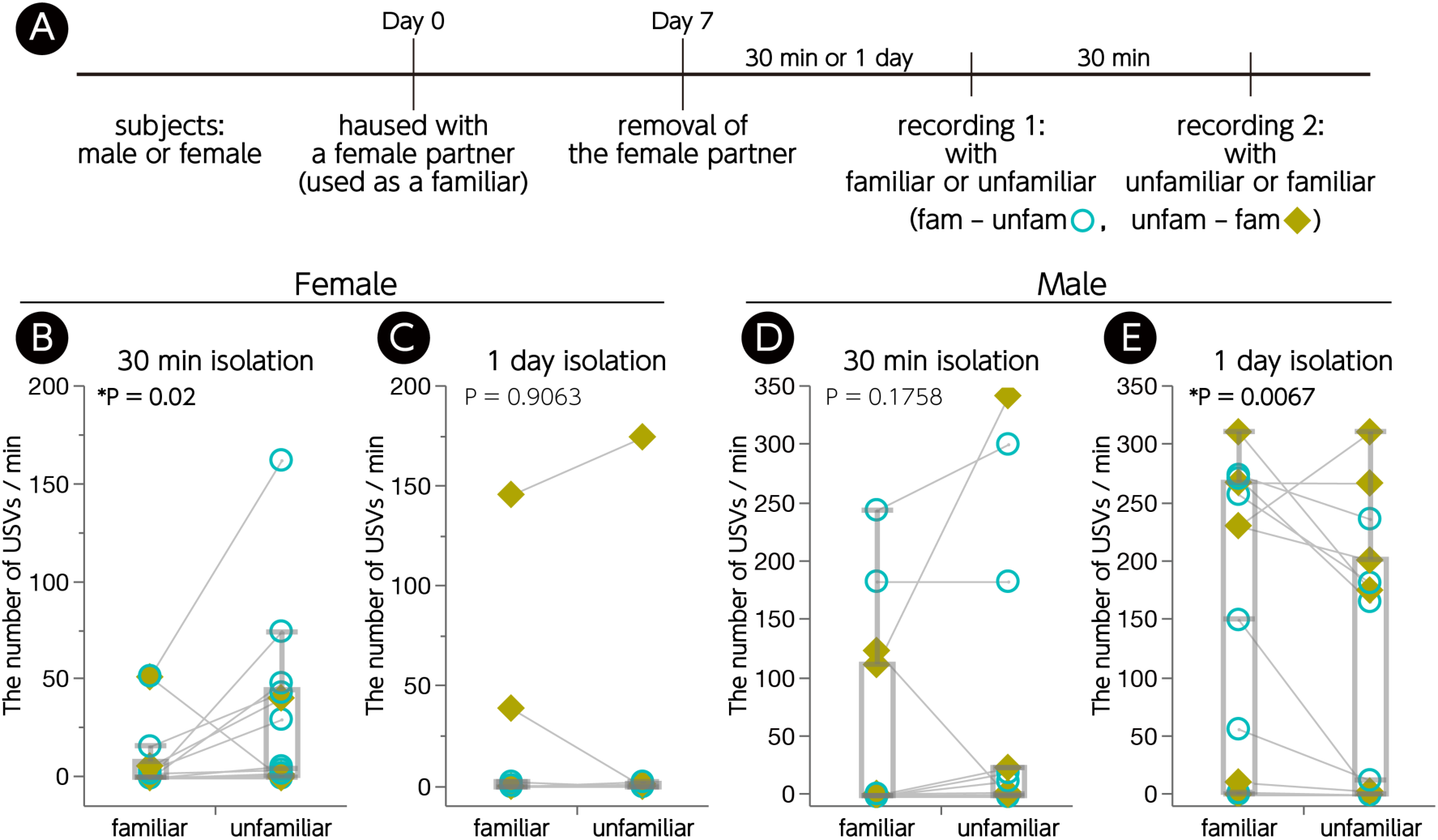
Male and female USVs to familiar or unfamiliar females. (A) Schematic diagram of the experimental procedure. (B) Female USVs to the familiar or unfamiliar females under the condition of 30 min isolation before recording 1. (C) Females’ USVs to the familiar or unfamiliar females under the condition of 1 day isolation before recording 1. (D) Male USVs to the familiar or unfamiliar females under the condition of 30 min isolation before recording 1. (E) Male USVs to familiar or unfamiliar females under the condition of 1 day isolation before recording 1. \**P* < 0.05, Wilcoxon signed-rank test; B, n = 11; C, n = 14, D and E, n = 15. Counter-balanced order of experimental contents for recording 1 and 2 in each individual is indicated as follows: Blue open circle, recording 1 with familiar and recording 2 with unfamiliar female (fam–unfam); lime yellow closed square, recording 1 with unfamiliar and recording 2 with familiar female (unfam–fam).

For both recordings, age-matched, unfamiliar adult females were also purchased, and 3–4 animals were housed in the same cage accordingly. These animals were used repeatedly as an unfamiliar intruder for recording (see below and Fig. 1A).

Food and water were supplied *ad libitum*, and the animals were kept under a standard 12-h:12-h light-dark cycle. All experiments were conducted in the light phase. The environment was maintained at a constant temperature (22–25°C) and humidity (50 ± 5%). All experimental procedures were approved by the Institutional Animal Use Committee of Kagoshima University (#L18004 and #L19003).

### 2.2 Experimental design and general procedures

The subject male was cohoused with an unfamiliar female, and the subject female was cohoused with a female that was bred together with the subject female in the same cage after weaning, as described above. These male–female and female–female mice were cohoused for 7 days without exchange of beddings. On the 7^th^ day (Day 7 shown in Fig. 1A) after the pairing of male–female and female–female began (Day 0 shown in Fig. 1A), the female partner was removed. Recordings of USVs were conducted twice (recording 1 and 2) for all subjects, and two experimental conditions were tested in both male–female and female–female contexts. Some subjects were used for recording 1 thirty min after the removal of the partner, and the other subjects were used 1 day (24–25 h) after the removal. Recording 2 was conducted 30 minutes after recording 1 for all subjects. USVs were induced by introducing a female mouse. The subjects were exposed to the familiar partner and an unfamiliar female once each, and the order of encountering the familiar or unfamiliar partner was counter-balanced between recordings 1 and 2.

### 2.3 Settings and procedures for ultrasound recordings

The microphone was hung 16 cm above the floor in a sound-attenuating chamber. Immediately prior to recording, the home cage of the subjects was moved into the chamber, where a red dim light was placed. The ultrasound recording was performed for 100 s after the intruder was introduced into the home cage.

According to previous studies (Ey et al., 2013; Ferhat et al., 2015; Matsumoto and Okanoya, 2016), we conducted recordings without the devocalization of female encounters under natural conditions. Previous experiments suggested that most vocalizations are produced by the resident subjects and that the vocal contribution of female encounters is very limited in the male–female and female–female contexts (White et al., 1998; Hammerschmidt et al., 2012; Neunuebel et al., 2015).

### 2.4 Ultrasound analysis

Most of the settings for the USV recordings followed those of our previous study (Kanno and Kikusui, 2018), with some modifications. Ultrasonic vocalizations were recorded using a CM16/CMPA condenser microphone (Avisoft Bioacoustics, Berlin, Germany), ultrahigh-speed ADDA converter BSA768AD-KUKK1710 and its software SpectoLibellus2D (Katou Acoustics Consultant Office, Kanagawa, Japan) with a sampling rate of 384 kHz (to measure 20-192 kHz).

Recorded sounds were saved on a computer as wav files using SpectoLibellus2D These sound files were analyzed with the GUI-based software USVSEG (implemented as MATLAB scripts) recently developed by us (Tachibana et al., 2020). Continuous sound signals with frequencies ranging from to 40160 kHz and durations ranging from 3 to 300 ms were analyzed and detected as syllables. Each syllable was segmented and saved as a JPEG image file, and the noise data (false positives) were manually excluded. Finally, the number of vocalizations (syllables) was quantified accordingly.

### 2.5 Statistics

Statistical analyses were performed using JMP 14 software (SAS Institute, Cary, NC, USA). In the present study, most of the data seemed to not be distributed normally; therefore, nonparametric tests were used. To compare the two conditions within an individual, Wilcoxon signed-rank tests were applied (see Fig. 1), and Kruskal–Wallis tests were used for comparison among groups, followed by post-hoc Steel–Dwass tests (see Figs. 2 and 3). Values were reported as dot plots with boxplots. Differences with *P* < 0.05 were considered significant.

**Fig. 2.**
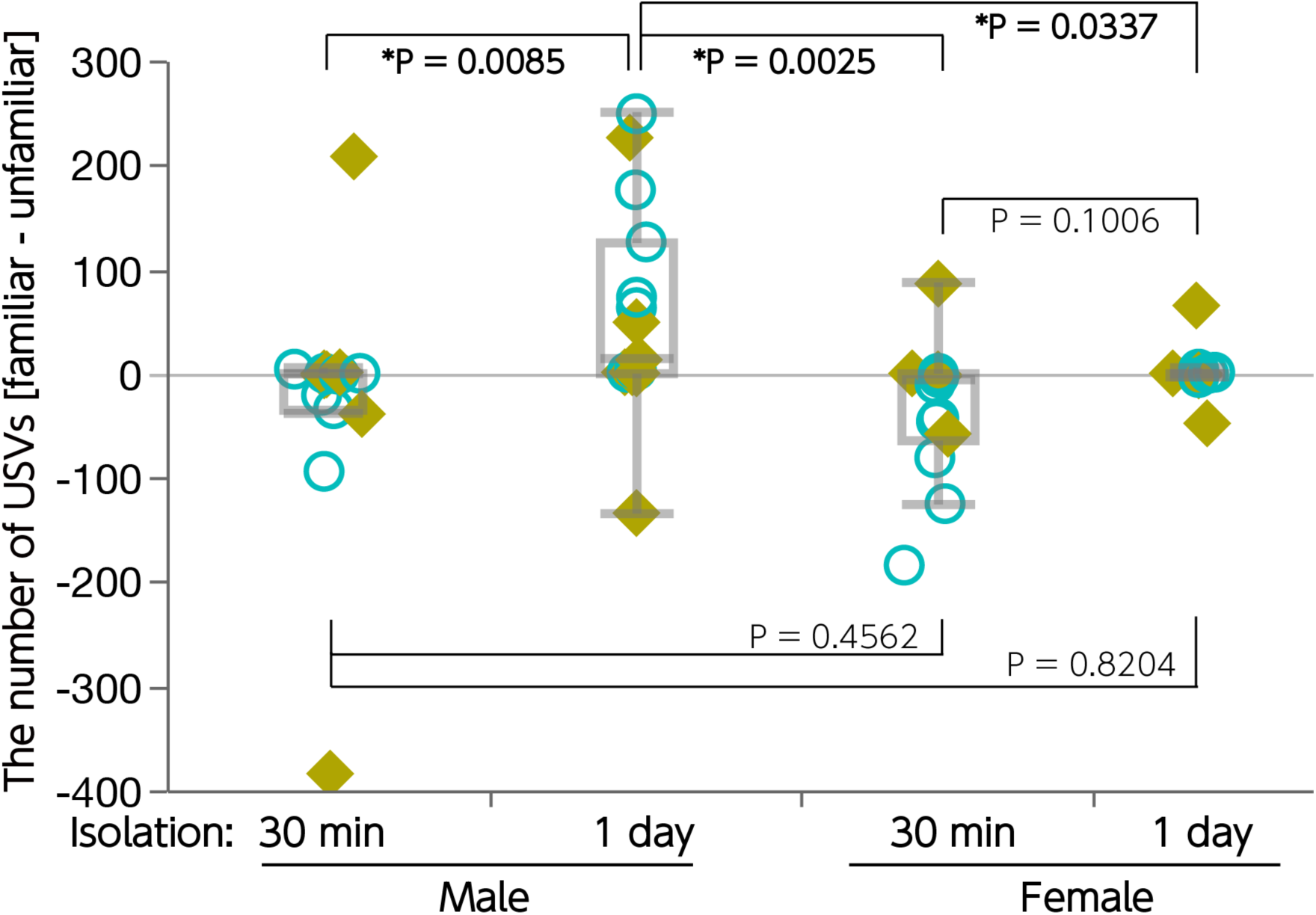
Sex differences in vocal response to familiar or unfamiliar females. A total comparison among groups is shown in Fig. 1. \**P* < 0.05, Steel–Dwass test after Kruskal–Wallis test; male 30 min and 1 day isolation, n = 15; female 30 min isolation, n = 11, female 1 day isolation, n = 14. The counter-balanced order of experimental contents for recording 1 and 2 in each individual is indicated in Fig. 1: Blue open circle, fam– unfam; lime yellow closed square, unfam–fam.

**Fig. 3.**
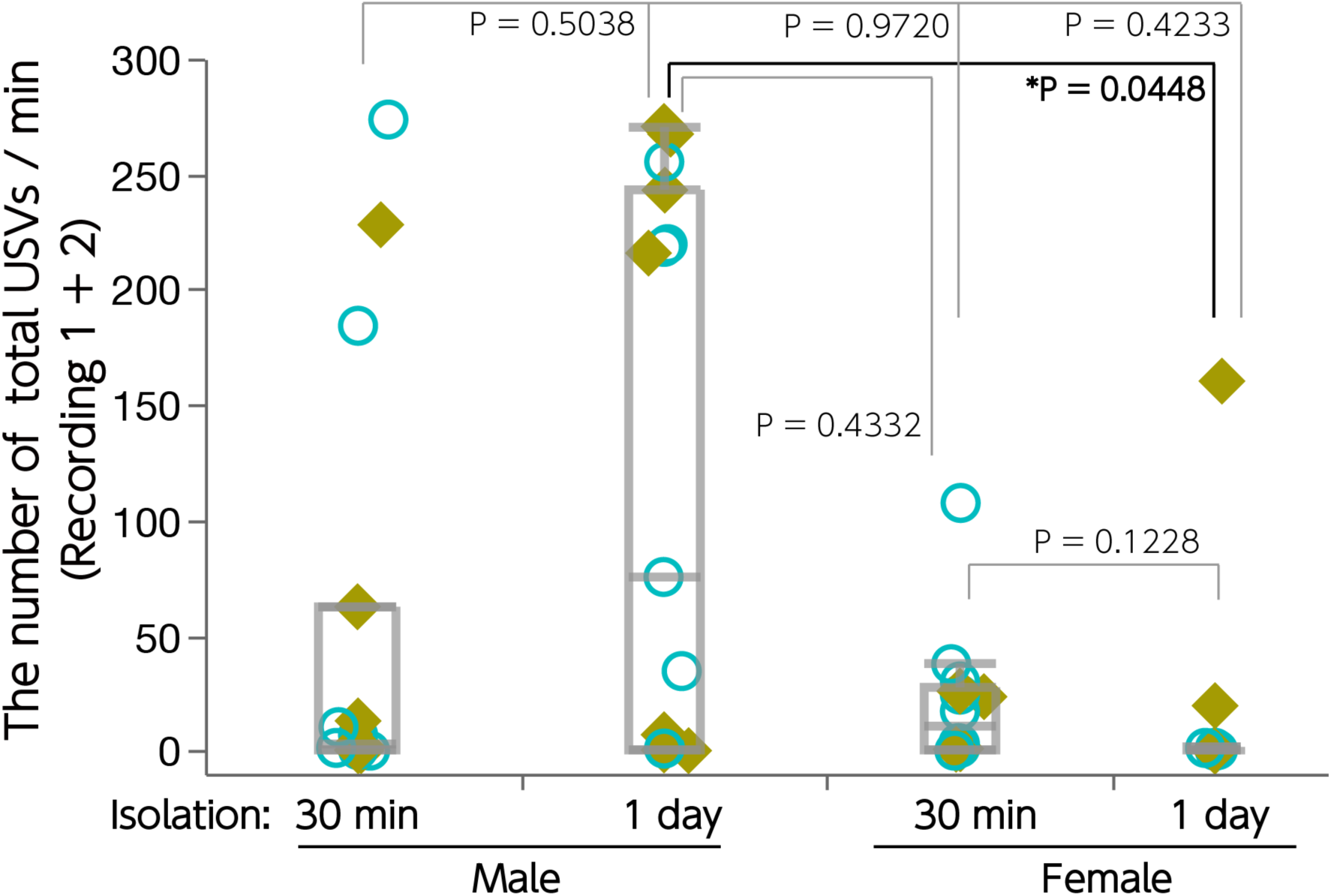
Comparison of the total number of USVs among groups. The total number of USVs (recording 1 + 2) per min among groups is shown. 1. \**P* < 0.05, Steel–Dwass test after Kruskal–Wallis test; male 30 min and 1 day isolation, n = 15; female 30 min isolation, n = 11, female 1 day isolation, n = 14. The counter-balanced order of experimental contents for recording 1 and 2 in each individual is indicated as in Fig. 1-2: Blue open circle, fam–unfam; lime yellow closed square, unfam–fam.

## 3. Results

### 3.1 Female–female vocalization

Under the 30 min isolation condition, the number of USVs emitted from subject females to unfamiliar females was significantly higher than that from familiar partners (Fig. 1B; Wilcoxon signed-rank test, *W*_(14, 14)_ = 39.0, *P* = 0.02). On the other hand, there was no significant difference between the number of USVs emitted to the unfamiliar and familiar partner after 1 day of isolation (Fig. 1C; Wilcoxon signed-rank test, *W*_(11, 11)_ = 3.0, *P* = 0.9063).

### 3.2 Male–female vocalization

Under the 30 min isolation condition, there was no significant difference between the number of USVs emitted from males to unfamiliar and familiar partners (Fig. 1D; Wilcoxon signed-rank test, *W*_(15, 15)_ = 26.0, *P* = 0.1758). On the other hand, the number of USVs emitted to the familiar partner was significantly higher than that of the unfamiliar female after 1 day of isolation (Fig. 1E; Wilcoxon signed-rank test, *W*_(15, 15)_ = −47.5, *P* = 0.0067).

### 3.3 Total comparison among groups

Differences between the number of USVs emitted to familiar and unfamiliar partners ([#. familiar] - [#. unfamiliar]) were calculated from the data shown in Fig. 1 and were compared among the 4 groups (Fig. 2). The Kruskal–Wallis test indicated significant differences between groups (χ^2^ = 19.4288, *df* = 3, *P* = 0.0002) and the Steel–Dwass test (performed *post-hoc*) showed that the value of males under the 1 day isolation condition was significantly higher than that of males or females under the 30 min isolation condition (*P* = 0.0085 and *P* = 0.0025, respectively), or females under the 1 day isolation condition (*P* = 0.0337).

Additionally, the total number of USVs were compared in the two recording tests (recording 1 + 2). The number of USVs in the 200-sec recording was calculated as USVs per minute in order to discuss the results with that of previous studies (Fig. 3). The Kruskal–Wallis test indicated significant differences between groups (χ^2^ = 9.0411, *df* = 3, *P* = 0.0287) and the Steel–Dwass test (performed *post-hoc*) showed that the value of males under the 1 day isolation condition was significantly higher than that of females under the 1 day isolation condition (*P* = 0.0448) respectively. No significant difference was observed in the other pairwise comparison of *post-hoc* tests.

## 4. Discussion

### 4.1 Female–female vocalization

As described in the Introduction, we prepared female subjects who spent more time with a female partner. Here, relations between them are thought to be affiliative. Nevertheless, contrary to our expectation, the result of the previous study of D’Amat and Moles (D’Amato and Moles, 2001) was replicated in our 30 min isolation condition (Fig. 1B). Here, we found that females vocalized more to an unfamiliar female than to familiar females. It is unclear why females rarely vocalized under the 1 day isolation condition, but no difference was found between the number of USVs to the unfamiliar and familiar females under such conditions, as was the case in D’Amat and Moles’ studies. We believe hence, that it is safe to conclude that B6 females vocalize more to unfamiliar females.

### 4.2 Male–female vocalization

On the other hand, it was surprising that males exhibited an opposite tendency toward females. It has been reported that male courtship USVs are enhanced by sociosexual experience (Kanno and Kikusui, 2018) and that vocal usage is altered according to the progressive phase of sexual behavior (Matsumoto and Okanoya, 2016). Furthermore, castration reduces the USVs, and replacement of sex hormones restores the vocalizations (Nunez et al., 1978). In particular, the implantation of testosterone in both the ventral tegmental area and medial preoptic area, which is known as a neural circuit for sexual motivation, effectively restores the USVs in castrated male mice (Sipos and Nyby, 1996b). These results indicate that male USVs directed towards females express sexual motivation as Nunez et al. suggested that “measures of male vocalizations provide an index of sexual motivation independent of male copulatory performance” (Nunez et al., 1978). We expected males to show more vocalization to unfamiliar females; however, contrary to our expectation, males made many vocalizations to the familiar partner. Even though different experimental conditions, such as length of cohousing, could possibly have led to different results, it appears that males exhibited more sexual motivation toward the familiar partner (cohoused for 1 week) under the present conditions.

### 4.3 Technical limitations and considerations for experimental design

It is not clear why the optimal conditions for observation of differential vocal responses to the familiar or unfamiliar females differ between males and females. Additionally, there was a potential design flaw in the present paradigm. A major difference between the female–female pairs and the male–female pairs is that the female– female pairs were housed together since the time of weaning, whereas the male–female pairs were housed together only for 7 days. This makes interpretation of any sex differences difficult. Nevertheless, sex differences in such vocal responses are still considered important. For decades, the sex differences between male–female and female– female USVs have been discussed. The amount of USVs between male–female and female–female is comparable (Maggio and Whitney, 1985). In addition, the vocal repertoire between sexes has also been reported to be comparable (Hammerschmidt et al., 2012). In contrast, the sex differences shown in the present study indicate that mice change their vocal behavior in response to the context of whom they are dealing; their perception of the context is based on social cognition and memory. As observed in the female vocal response —this trend in response was still reproduced in the present study under different conditions from a previous study, mice usually exhibit attention to novel things or individuals. In some behavioral tests, such as habituation–dishabituation test, this property is being utilized accordingly. However, because males vocalized more to familiar females, USVs are thought to be not just a response to novel things or individuals but also to be a social response. These findings with wild-type B6 mice provide a practical behavioral assay for the disease model mice, such as genetically modified mice with B6 background, focusing on their social impairment. This is because if typical social behavior can be observed in the wild-type both as a quality as well as a quantity, then a valid assay for detection of atypical characteristics as a deviation from *normal* in such a disease model can be established accordingly.

Another issue is that the paired experimental design seems problematic in combination with the fact that there was a 30 min delay between exposure to the familiar and unfamiliar female; the memory retention period was 30 min for the first exposure and 60 min for the second exposure. Therefore, though some would argue that between-comparison would have been better, there is yet another concern. There are large individual differences, at least in male–female USVs (Kanno and Kikusui, 2018). In fact, we conducted a Wilcoxon signed-rank test using only the data of recording 1 (therefore, N values became half) and compared the effect of familiarity of encountering females as a between-factor; however, no significant difference was observed in any group. In such cases, a better way to observe the effects of the experiment would be to take repeated measurements while taking individual differences into account. Notably, the present study demonstrated such significant differences using a within-experimental design. In addition, since differential vocal response in subject females to familiar or unfamiliar ones was significantly observed with 30- and 60-min retention times, as mentioned in the Introduction (D’Amato and Moles, 2001), we speculate that our present paradigm of repeated measurements could detect such difference before extinction of memory for social partners.

Again, it is unclear why females rarely vocalized under the 1 day isolation condition. Similarly, depending on the isolation conditions, the amount of USVs seems different in males. This might be explained by sexual refractory periods in males since they do not exhibit sexual behaviors (mounting and intromissions) after ejaculation. This interval between sexual behaviors is called a refractory period and USVs are not observed in this period. (Nyby, 1983). In the present study, males vocalized more under the 1 day isolation than under the 30 min isolation condition, possibly because males were thought to be sexually saturated as a result of the 7 days of cohousing with a female before experiments. This 1 day isolation condition was thought to restore their sexual motivation and was tested because we expected this possibility. Nevertheless, it must be noted that the observed USVs in this study are low not only with respect to females but also for the males. The mean and median of USVs / min in this study (Fig. 3) are as follows ([mean, median]): male 30 min isolation [52.3, 3]; male 1 day isolation [120.6, 75.3]; female 30 min isolation [19.7, 10.2]; female 1 day isolation [16.7, 0.3]. On the other hand, the number of USVs/min reported in other studies, including our previous study, is higher. For example, the mean number of approximately 250 / min (Kikusui et al., 2011; Kanno and Kikusui, 2018; Hori et al., 2020) or approximately 100-150 / min (Hammerschmidt et al., 2012; Ey et al., 2013) of male–female USVs, and approximately 150-200 / min of female–female USVs have also been reported (Hammerschmidt et al., 2012; Ey et al., 2013). In these studies, several days (up to 7 days) of single housing was carried out in both male and female subjects. This is because sexual motivation is enhanced — and this motivation leads to increased USVs— for males, as mentioned above. In addition, Ey et al. explained “Tested females were isolated 3 days before the resident–intruder test to increase their motivation for social interactions” (Ey et al., 2013). Thus, in general, several days of isolation are apt to enhance USVs, even though long-term isolation cannot be used before recording to retain the memory of a social partner. Therefore, such studies focusing on social relations that are formed in the process of relatively long-term social interaction must be limited in their experimental paradigm. Even under such restrictions, we reported new findings in this study. For example, though the reason or mechanisms of isolation condition affecting the amount of USVs was identified, significant differences observed under certain conditions were dependent on differences in social partners (familiar or unfamiliar).

### 4.4 Functional consideration of female–female USVs

Although such sex differences in USVs have been revealed, the past and present studies have yet to determine the biological significance and function of female–female USVs. Since Maggio and Whitney (Maggio and Whitney, 1985) hypothesized that female vocalization contributes to the formation of social hierarchy among females, this hypothesis has been referred to in some studies (D’Amato and Moles, 2001; Hammerschmidt et al., 2012). However, to our knowledge, this hypothesis has never been investigated experimentally in mice. A recent study made an important point about sex differences in acoustic features, with more diversity (inter-individual difference) observed in the vocal repertoire in males than in females (Matsumoto and Okanoya, 2018). Nonetheless, the biological significance and function of female USVs remains unclear. Under such circumstances, we could speculate, at least a possible role based on social behaviors as already observed among females. It has been reported that female house mice display nonrandom preferences to other females, that is, a social partner choice. The partner choice in pairs with a preferred one leads to a higher probability of giving birth and establishing a cooperative relationship for communal nursing of offspring, resulting in higher reproductive success (Weidt et al., 2008). Female–female USVs may function in the process of such partner choice; however, further studies focusing on clarifying roles of the USVs, including investigation of the associated neural mechanisms, are needed.

## 5. Conclusion

We found that females vocalized more to unfamiliar than to familiar females in B6 strains, which could be replication as observed in another strain. In contrast, males exhibited more USVs to familiar partners. This sexually dimorphic behavior suggests that mice change their vocal behavior in response to the social context, and that their perception of the context is based on social cognition and memory. In addition, because males vocalized more to familiar females, USVs appear to be not just a response to novel things or individuals, but also could be a social response. The mechanisms and roles of these behaviors should be investigated further; however, these behaviors can be useful assays for social behavior to compare with some transgenic strains, which can be applied to both males and females, respectively.

## Ethics

All experimental procedures were approved by the Institutional Animal Use Committee of Kagoshima University (#L18004 and #L19003).

## Authors’ contributions

K.K. contributed to the experimental design. E.S. and Y.T. conducted the experiments. E.S., Y.T., and K.K. contributed to the interpretation of data and writing of the manuscript.

## Acknowledgments

This work was supported by a Grant-in-Aid for Scientific Research on Innovation Areas “Integrative Research toward Elucidation of Generative Brain Systems for Individuality” (19H04912) from MEXT to K.K. We thank various supports, including daily discussion, of this project. We would like to thank Editage (www.editage.com) for English language editing.

